# Identification of germ cell-specific *Mga* variant mRNA that promotes meiotic entry via impediment of a non-canonical PRC1

**DOI:** 10.1101/2020.09.03.282368

**Authors:** Yuka Kitamura, Kousuke Uranishi, Masataka Hirasaki, Masazumi Nishimoto, Ayumu Suzuki, Akihiko Okuda

## Abstract

Transition from mitosis to meiosis in cell division is a fundamental process of gametogenesis. This transition is thought to be largely controlled by the exchange of relative dominance between positive and negative regulation by the retinoic acid/Stra8 signal cascade and a non-canonical PRC1 (PRC1.6), respectively. We have previously demonstrated that germ cells have transcriptionally and/or post-translationally reduced levels of MAX, a component of PRC1.6, immediately prior to meiotic onset, leading to alleviation of the negative effect of PRC1.6 against meiotic onset. Here, we found that germ cells produced *Mga* variant mRNA bearing a premature termination codon (PTC) during meiosis as an additional mechanism to impede the function of PRC1.6. Our data indicated that spermatocytes and/or round spermatids produced an anomalous MGA protein lacking the bHLHZ domain from the variant mRNA and therefore functioned as a dominant negative regulator of PRC1.6 by exquisitely using their inefficient background of PTC-mediated nonsense-mediated mRNA decay. Thus, our data indicate that meiotic onset of male germ cells is controlled in a multi-layered manner in which both MAX and MGA, which constitute the core of PRC1.6 by their interaction, are at least used as targets to deteriorate the integrity of the complex to ensure initiation of meiosis.

**Significance Statement:** PRC1.6, a non-canonical PRC1, functions as a strong blocker of meiotic onset. Therefore, germ cells need to alleviate the function of the complex as a prerequisite for meiotic onset. The MGA/MAX heterodimer not only constitutes a core of PRC1.6, but also confers direct DNA-binding activity to the complex. We have previously demonstrated that germ cells reduce Max amounts prior to meiotic onset to inactivate PRC1.6. In this study, we explored the possibility of an additional molecular mechanism that promotes meiotic onset via impediment of PRC1.6 functions as a safeguard system. Here, we demonstrate that meiotic germ cells specifically generate variant *Mga* mRNA by alternative splicing, which leads to production of a dominant negative regulator of PRC1.6.

## Introduction

Meiosis is a specialized type of cell division, which converts a cell from diploid to haploid (1). Signaled by retinoic acid 8 (Stra8) is a crucial positive regulator of meiosis in mammals and molecular cloning of Meiosin, which functions by forming a complex with Stra8, significantly advanced our understanding of the molecular bases of Stra8-dependent meiotic onset (2). In contrast to the STRA8/MEIOSIN complex, polycomb repressive complex 1 (PRC1) is a negative regulator of meiotic onset in mammals (3, 4). The PRC1 family includes six distinct subtypes, PRC1.1–6, which share RING1A or RING1B as a common subunit bearing enzymatic activity for ubiquitination of histone H2A at lysine 119, but differ in their composition of other subunits (5–9). The PRC1 family is largely classified into two groups, i.e., canonical (PRC1.2 and PRC1.4) and non-canonical (PRC1.1, PRC1.3, PRC1.5, and PRC1.6) complexes. Canonical PRC1s containing chromobox proteins bind to chromatin via an interaction with the histone modification H3K27me3 catalyzed by PRC2. Therefore, recruitment of canonical PRC1s to chromatin occurs after binding of PRC2 (10–12). However, non-canonical PRC1s such as PRC1.6, which do not recognize this histone modification, are recruited to chromatin prior to binding of PRC2 using their distinct ways one another (5, 13, 14). For example, recruitment of PRC1.1 to genome target sites is dependent on its KDM2B subunit that recognizes non-methylated CpG islands (15). In terms of PRC1.6, two DNA-binding proteins of the complex, i.e., MGA and E2F6, are used for its direct binding to genomic sites (5, 16). It is also known that PCGF6, L3MBTL2, RYBP, and YAF2 substantially contribute to recruitment of PRC1.6 to chromatin (17, 18). We and others have recently demonstrated that PRC1.6 acts as a strong blocker of ectopic and precocious onset of meiosis in embryonic stem cells and germ cells, respectively (19–21), which suggests that a previous report by Yokobayashi et al. (3) showing strong induction of meiosis by deprivation of RING1B, a common component in the PRC1 family, is largely accounted for by disruption of PRC1.6. We have also demonstrated that germ cells transcriptionally and/or post-translationally reduce their amount of MAX, a component of PRC1.6, to liberate them from PRC1.6-dependent repression at the timing of or immediately prior to meiotic onset (19). However, because switching from mitosis to meiosis in cell division is of paramount importance for gametogenesis, we assumed that inactivation of PRC1.6 in germ cells is not solely dependent on the reduction of Max protein levels, but regulated in a multi-layered manner to ensure onset of meiosis as a safeguarding system.

Here, we addressed this issue by searching for potential exon sequences in genes encoding one of the components of PRC1.6 using SpliceAI, a recently developed deep neural network (22), which led to identification of *Mga* variant mRNA carrying a novel sequence with a premature termination codon (PTC). We also found that this variant mRNA generated by alternative splicing is specifically present in meiotic germ cells and round spermatids. Furthermore, our data demonstrated that meiotic spermatocytes and spermatids translate the variant mRNA into carboxy-terminally truncated MGA protein that functions as a dominant negative regulator of PRC1.6 owing to the lack of the basic helix-loop-helix/leucine zipper (bHLHZ) domain by taking advantage of the inefficient background of nonsense-mediated mRNA decay (NMD) in these cells.

## Results

### Exon inclusion is the most prevalent type of alternative splicing during transition from mitosis to meiosis in germ cells

We have previously demonstrated that germ cells physiologically reduce their amount of MAX protein that constitutes the core of PRC1.6 with MGA to de-repress meiosis-related genes prior to meiotic onset (19). In this study, we explored the possibility of an additional molecular mechanism that inactivates the function of PRC1.6 to ensure meiotic entry. Because the testis is known for its prevalence of alternative splicing similar to the brain (23, 24), we pursued the possibility of involvement of alternative splicing in facilitating meiosis by deteriorating the function of PRC1.6. First, we examined which stages in spermatogenesis and which types of alternative splicing were prevalent in germ cells by inspecting publicly reported RNA sequence data. These analyses revealed that alternative splicing occurred most actively during meiotic onset (transition of spermatogonia to preleptotene spermatocytes) (Fig. S1A). In terms of the types of alternative splicing, the skipping exon (SE) type was the most prevalent, which accounted for more than 50% of all alternative splicing events among the five distinct splicing types. Furthermore, our analyses revealed that gain of a novel exon was approximately twice as frequent as loss of an exon among the SE-type alternative splicing events (Fig. S1B). Moreover, our data revealed that most transcripts that gained or lost an exon around meiotic onset maintained their forms by *de novo* synthesis and/or stabilization at least up to the round spermatid stage (Fig. S1C). We also classified alternative splicing events in neural progenitor cells and mesenchymal stem cells using publicly reported RNA sequence data. These analyses revealed that SE was also the most prevalent alternative splicing during differentiation of neural progenitor cells and mesenchymal stem cells (Fig. 1SA). However, comparisons of genes that gained a new exon in germ cells with those in neural cells and mesenchymal stem cells revealed that these three gene populations barely overlapped (Fig. 1SD), which indicated that at least genes subjected to this type of alternative splicing were selected distinctly in each cell type.

### Identification of testis-specific *Mga* variant mRNA

Based on these data, we explored the possibility that the SE type of alternative splicing, particularly gain of a novel exon, was involved in the inactivation of PRC1.6 during the transition from mitosis to meiosis in germ cells. Because SpliceAI, a deep neural network that predicts mRNA splicing from a genomic sequence (22), was developed recently, we used this technology to identify putative exon sequences within genes encoding a component of PRC1.6. First, we confirmed that SpliceAI identified all exons of genes encoding a component of PRC1.6 (*Mga*, *Max, L3mbtl2, E2f6, Rnf2*, and *Pcgf6*) (Figs. 1A and S2), which validated the accuracy of prediction by this deep learning program. More importantly, SpliceAI additionally predicted six genomic regions as putative exons (one each within *E2f6* and *Pcgf6* genes and two each in *Mga* and *L3mbtl2* genes) (Figs. 1A and S2). Therefore, we performed RT-PCR analyses of RNAs from various tissues to determine the possibility that RNAs transcribed from these regions were incorporated as exon sequences into mature mRNA in certain tissues (Fig. S3). These analyses revealed that a putative exon located within the 18^th^ intron of the *Mga* gene was specifically incorporated into RNA from the testis, but only marginal presence was evident in other tissue RNAs. However, the other five regions were not entirely incorporated into any examined tissue RNAs. Because of the testis specificity, we further investigated alternative splicing using a portion of the 18^th^ intron of the *Mga* gene as an exon. First, we confirmed testis-specific incorporation of the transcript from this region by quantitative PCR (qPCR) (Fig. 1B, upper panel). According to the scores calculated by SpliceAI (0.695 and 0.761 for the splice acceptor and donor, respectively), this region was not predicted to be a constitutive exon (more than 0.9, usually close to 1.0), but as an exon subjected to a substantial degree of alternative splicing regulation (usually between 0.1 and 0.9). Thus, our finding that this region was used for testis-specific alternative splicing was compatible with the prediction of SpliceAI. Therefore, we termed this sequence and *Mga* variant mRNA carrying this sequence as exon 19a and *Mga* splice variant (SV), respectively. With respect to wildtype *Mga* mRNA, we confirmed broad expression in various cell types. We also noted its relatively higher expression in the testis compared with any other examined tissues (Fig. 1B, lower panel).

**Figure 1.**
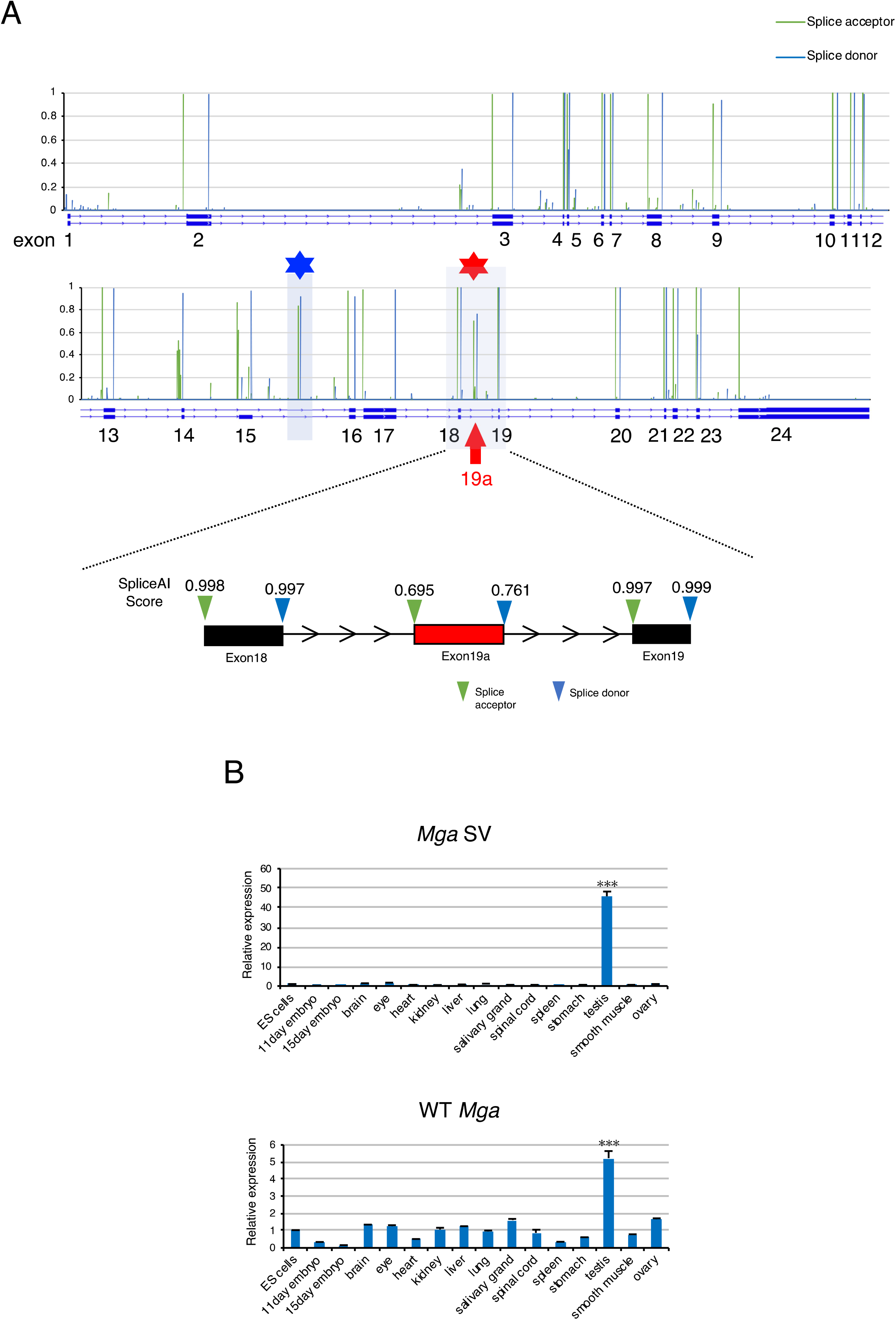
Identification of a novel testis-specific *Mga* splice variant. (A) Prediction scores as the splice acceptor and donor of *Mga* pre-mRNA from SpliceAI deep learning. Scores as the splice acceptor and donor are shown as green and blue bars, respectively, whose height is proportional to the score level. In addition to known exons, SpliceAI indicated a set of high scores for the splice acceptor and donor within the regions of the 15^th^ and 18^th^ introns of the *Mga* gene marked with blue and red asterisks, respectively. The latter region is enlarged to provide actual scores from SpliceAI. (B) qPCR analyses of *Mga* SV (upper panel) and wildtype *Mga* (lower panel) mRNAs in total RNAs from various tissues. Values obtained from ESCs were arbitrarily set to one for both *Mga* mRNA species. Data represents the mean ± standard deviation of three independent experiments. The Student’s t-test was conducted to examine statistical significance. \*\*\**P*<0.001

### RNA *in situ* hybridization analyses of *Mga* SV expression in the testis and epididymis

To determine which portions of the testis expressed *Mga* SV, RNA *in situ* hybridization analyses were performed using testis and epididymis tissues of adult mice. To avoid the difficulty associated with the short length (145 bp) of the *Mga* SV-specific sequence, we employed the BaseScope *in situ* hybridization technique rather than the conventional RNA hybridization method (25). These analyses clearly demonstrated that *Mga* SV-positive cells were not present in the interstitium portion, but restrictively present within seminiferous tubules, while wildtype *Mga*-positive cells were detected in both portions (Fig. 2A). These analyses also revealed that cells positive for wildtype *Mga* and those for *Mga* SV were abundantly and scarcely detected in the epididymis, respectively (Fig. 2B).

**Figure 2.**
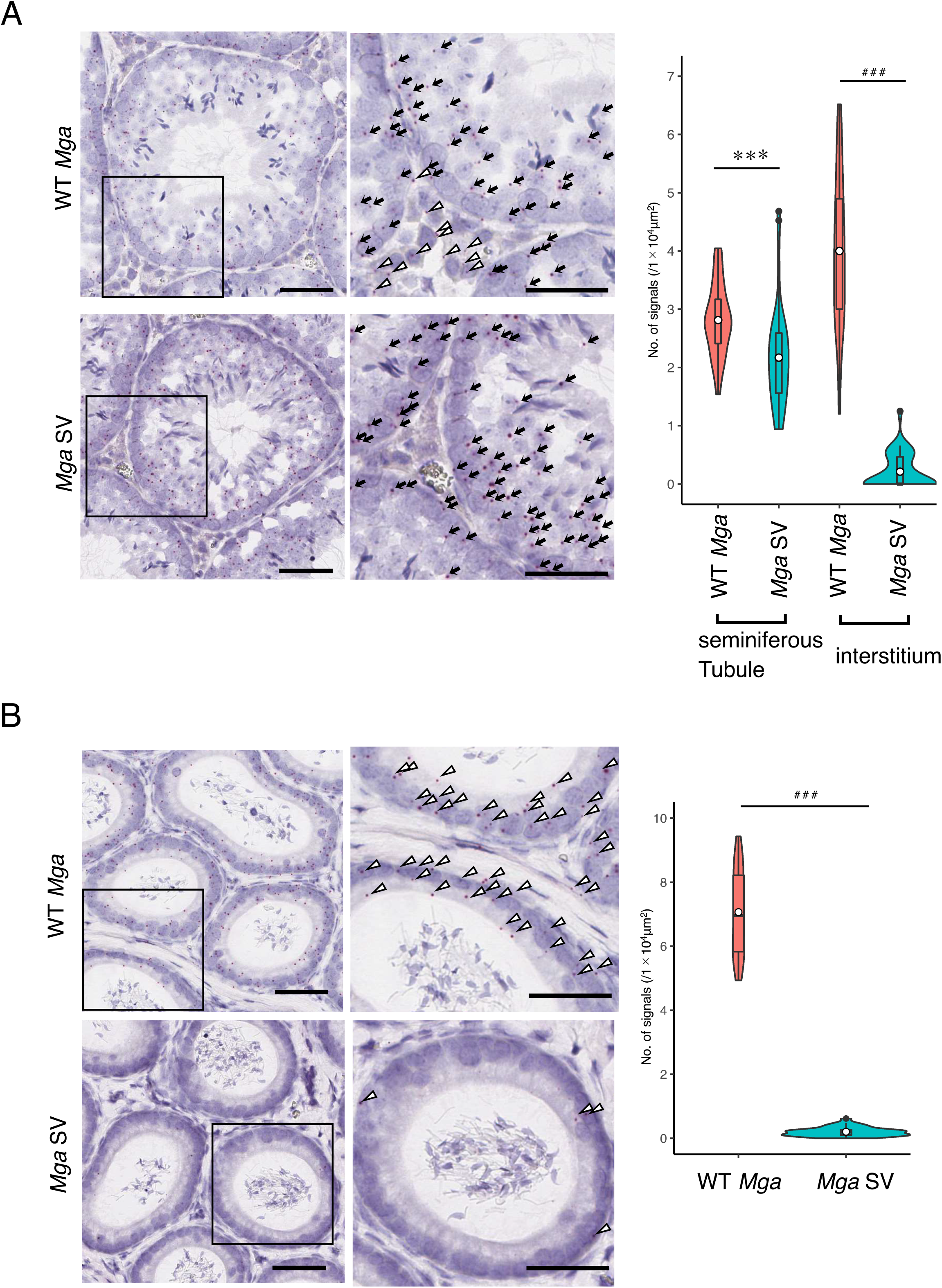
RNA *in situ* hybridization analyses of wildtype and variant *Mga* mRNAs in the testis and epididymis. (A) Detection of wildtype and variant *Mga* mRNAs in adult mouse testis by the BaseScope *in situ* hybridization technique. The region indicated with an open square is enlarged and shown at the right. Black arrow and open arrowhead indicate transcripts detected within seminiferous tubule (ST) and interstitium (I) portions, respectively. Data obtained as the numbers of positive signals in 35 randomly selected areas (0.1 mm^2^) were used to construct violin plots. To examine statistical significance, F-test values were first obtained for the respective data. Then, the Student’s t-test was conducted when the F-test value was larger than 0.05, whereas Welch’s t-test was conducted when its value was smaller than 0.05. \*\*\**P*<0.001 (Student’s t-test); ^###^*P*< 0.001 (Welch’s t-test) (B) Detection of wildtype and variant *Mga* mRNAs in the epididymis by the BaseScope *in situ* hybridization technique. Detection of wildtype *Mga* mRNA and its splice variant in the adult mouse epididymis and construction of violin plots as described in A. Region indicated with an open square is enlarged and shown at the right. Specific signals for wildtype and variant *Mga* mRNAs indicated by the open arrowhead are shown in the upper and lower panels, respectively. Data were analyzed statistically as described in A. ^###^*P*< 0.001 (Welch’s t-test)

### Production of *Mga* SV is coupled to meiosis of both male and female germ cells

To determine whether restrictive expression of *Mga* SV in cells within seminiferous tubules of the testis represented specific expression in germ cells, we prepared four distinct germ cell types, i.e., undifferentiated spermatogonia (Thy1^+^) and differentiated spermatogonia (c-Kit^+^) from testes of mouse pups at postnatal days 5–8 and spermatocytes and round spermatids from adult mice testes. Then, qPCR analyses were conducted using total RNAs from these cell populations. The analyses revealed that the levels of wildtype *Mga* mRNA were comparable among the four cell populations, whereas much higher expression of *Mga* SV was evident in spermatocytes and spermatids compared with Thy1^+^ and c-Kit^+^ spermatogonia in which meiotic initiation is blocked (26) (Fig. 3A). These results indicated that germ cells in the testis generated *Mga* SV and the timing of generation was coincidental with the timing of transition from mitosis to meiosis in cell division. Next, we examined publicly reported RNA sequence data of this sequence in spermatogonia, round spermatids, and germ cells undergoing meiosis in the testis (preleptotene and pachytene spermatocytes). In accordance with our qPCR data (Fig. 3A), visualization of publicly reported RNA sequence data of this putative exon (exon 19a) by Sashimi plot revealed specific production of *Mga* SV in meiotic spermatocytes and round spermatids, but not in spermatogonia (Fig. 3B). Male germ cells initiate meiosis in the testis at the beginning of puberty, whereas female primordial germ cells (PGCs) undergo meiosis and proceed to the diplotene stage of meiotic prophase I in the gonads during the mid-gestation stage (27, 28). Therefore, we inspected the publicly reported RNA sequence data to determine whether female PGC-specific onset of meiosis in the embryonic stage was also accompanied by the production of *Mga* SV. The analyses revealed that expression of *Mga* SV in female PGCs of the gonads became detectable from 11.5 to 16.5 dpc with peak expression at 15.5 dpc when meiotic germ cells are mostly at the pachytene stage, whereas no apparent peak for the production of *Mga* SV was detected in male germ cells at embryonic stage (Fig. 3C). Taken together, our data demonstrated an intimate link in the time scale between expression of *Mga* SV and meiosis in both male and female germ cells. Because female PGCs undergo meiosis at around 13 dpc, initiation of *Mga* SV production appeared to occur immediately prior to meiotic onset. It is also noteworthy that exon 19a bears in-frame stop codon (Fig. 3D), indicating that the coding sequence located downstream of the stop codon of exon 19a sequence, including that for the bHLHZ domain, was nullified. Intriguingly, inspection of the publicly reported RNA sequence data revealed that insertion of a PTC in conjunction with gain of a new exon was approximately three times more frequent in the transition from spermatogonia to germ cells at the preleptotene stage compared with differentiation of neural progenitor cells (Fig. 3E). These results indicated that production of *Mga* SV with a PTC represented typical alternative splicing of new exon inclusion in meiotic germ cells.

**Figure 3.**
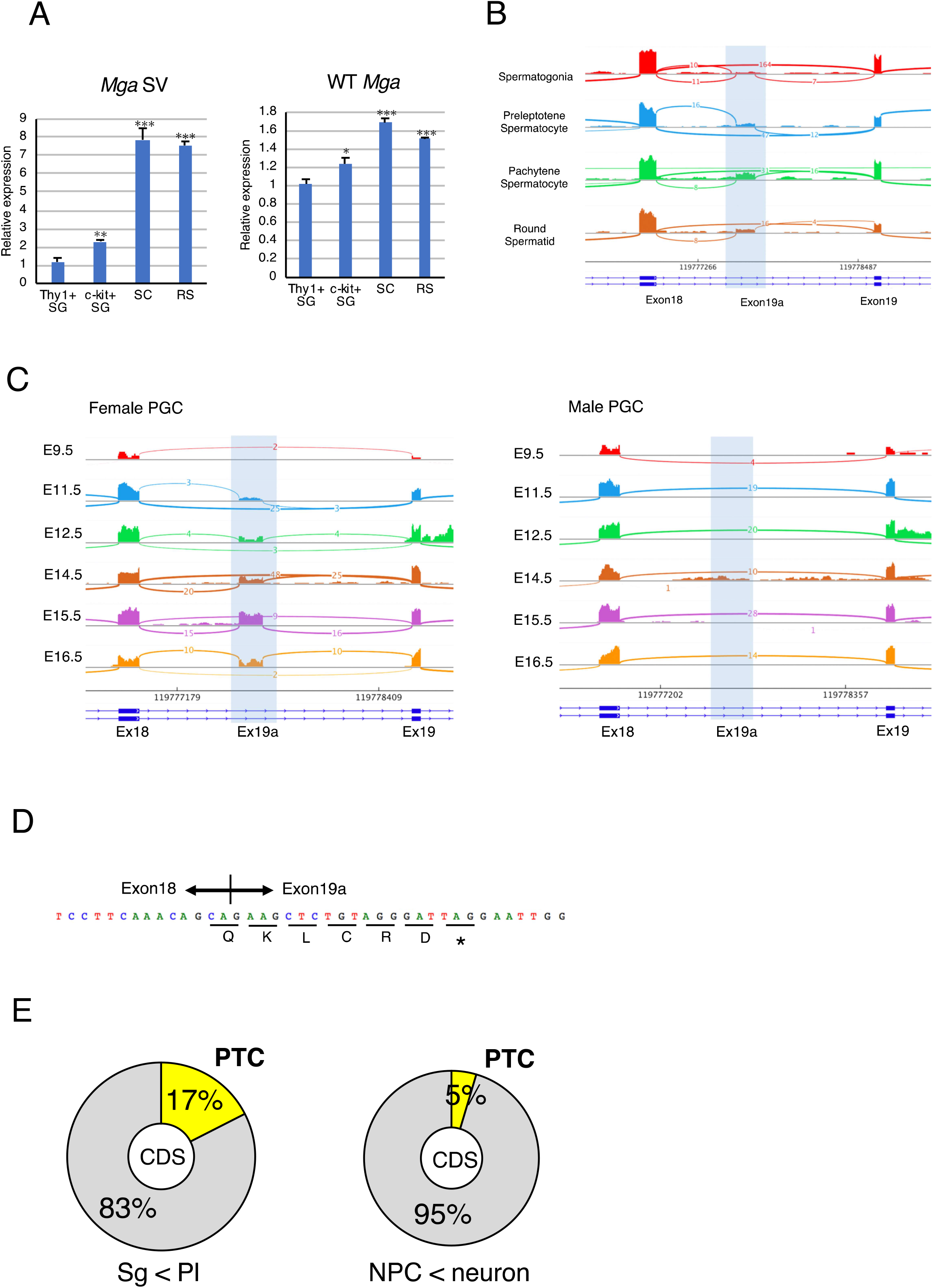
*Mga* splice variant is restrictively present around the meiotic stage of germ cells. (A) qPCR analyses of RNAs from undifferentiated spermatogonia (Thy1^+^), differentiated spermatogonia (c-Kit^+^), spermatocytes, and round spermatids to determine the levels of *Mga* SV (left panel) and wildtype *Mga* (right panel) mRNAs. Values obtained from Thy1^+^ spermatogonia were arbitrarily set to one for both *Mga* mRNA species. Data were subjected to statistical analyses as described in Figure 1B. \*\**P*<0.01; \*\*\**P*<0.001 SC and RS indicate spermatocytes and round spermatids, respectively. (B) Visualization of splicing events around exons 19 and 20 of the *Mga* gene during spermatogenesis in the testis by Sashimi plots. Plots were constructed using publicly reported data (GSE75826). Abbreviations are described in A. (C) Sashimi plots showing splicing events around exons 19 and 20 of the *Mga* gene in male and female PGCs during the embryonic stage. Data for female and male PGCs were obtained from publicly reported data (E-MTAB-4616) and shown in left and right panels, respectively. (D) Sequence of exon 19a containing the in-frame PTC. Acquisition of the exon 19a sequence by alternative splicing was accompanied by insertion of PTC after the addition of the coding sequence of five amino acids, leading to translation into the carboxy-terminally truncated anomalous MGA protein that lacked the bHLHZ domain. (E) Generation of the PTC-containing transcript by alternative splicing was a frequent event during meiotic onset of germ cells. Frequencies of exon inclusion leading to incorporation of a PTC and that leading to addition of the novel amino acid sequence occurred during conversion of spermatogonia to meiotic germ cells (left) and differentiation of neural progenitor cells into neuronal cells (right) are presented as pie charts. Publicly reported RNA sequence data of germ cells (GSE75826) and neuronal cells (GSE96950) were used to calculate the frequencies.

### Inefficiency of NMD substantially contributes to accumulation of *Mga* SV transcripts in meiotic spermatocytes and spermatids

Notably, an in-frame terminating codon was present in exon 19a (Fig. 3D). In general, transcripts containing a PTC are subjected to rapid degradation by NMD (29–31). However, meiotic and post-meiotic germ cells in the testis are defective for PTC-mediated NMD, but not long 3’-untranslated region (UTR)-mediated NMD (32, 33). Therefore, we speculated that an inefficient background of PTC-mediated NMD substantially contributed to the specific presence of the *Mga* SV transcript in meiotic spermatocytes and round spermatids. To test this hypothesis, three germ cell populations [germline stem cells (GSCs), spermatocytes (SCs), and round spermatids (RSs)] and two types of non-germ cells [mouse embryonic fibroblasts (MEFs) and embryonic stem cells (ESCs)] were treated with cycloheximide (CHX) that inhibits NMD. Then, RNAs of these cells were used to quantify expression levels of *Mga* SV by semi-quantitative and quantitative PCRs after conversion to cDNAs (Fig. 4A and B). These analyses revealed that CHX treatment clearly augmented the amounts of *Mga* SV transcripts in ESCs, MEFs, and GSCs compared with untreated cells, which indicated that *Mga* SV mRNA produced in these cells was indeed subjected to degradation by NMD. However, no noticeable alterations in the amounts of *Mga* SV due to treatment with CHX were observed in spermatocytes and round spermatids, which was consistent with the notion of low PTC-mediated NMD activity in these cells. We also noted that the levels of *Mga* SV mRNAs in CHX-treated ESCs and MEFs were significantly lower than those in spermatocytes and round spermatids, whereas CHX-treated GSCs showed almost equivalent levels of *Mga* SV mRNAs as those in untreated spermatocytes and spermatids. Therefore, these results indicated that specific accumulation of *Mga* SV mRNAs in spermatocytes and round spermatids represented the combined consequence of two independent phenomena, i.e., preferential production of *Mga* SV mRNA in germ cells and attenuated NMD activity against PTC-containing mRNAs in meiotic germ cells. Next, we examined whether the presence of *Mga* SV mRNAs in spermatocytes and spermatids was associated with production of carboxy-terminally truncated MGA protein. To this end, we prepared nuclear extracts from ESCs and a mixture of spermatocytes and round spermatids and then performed western blot analyses. The analyses using an anti-MGA antibody that recognized the amino-terminal portion of MGA detected a specific band corresponding to the expected size of the protein translated from the initiating methionine to PTC of *Mga* SV mRNA (hereafter MGA SV) as well as a band corresponding to wildtype MGA, whereas only a band corresponding to wildtype MGA was detected in the protein extract from ESCs (Fig. 4C, left panel). The lack of the carboxy-terminal portion of MGA SV was confirmed by analyses using an antibody that recognized the carboxy-terminal portion of MGA (Fig. 4C, right panel).

**Figure 4.**
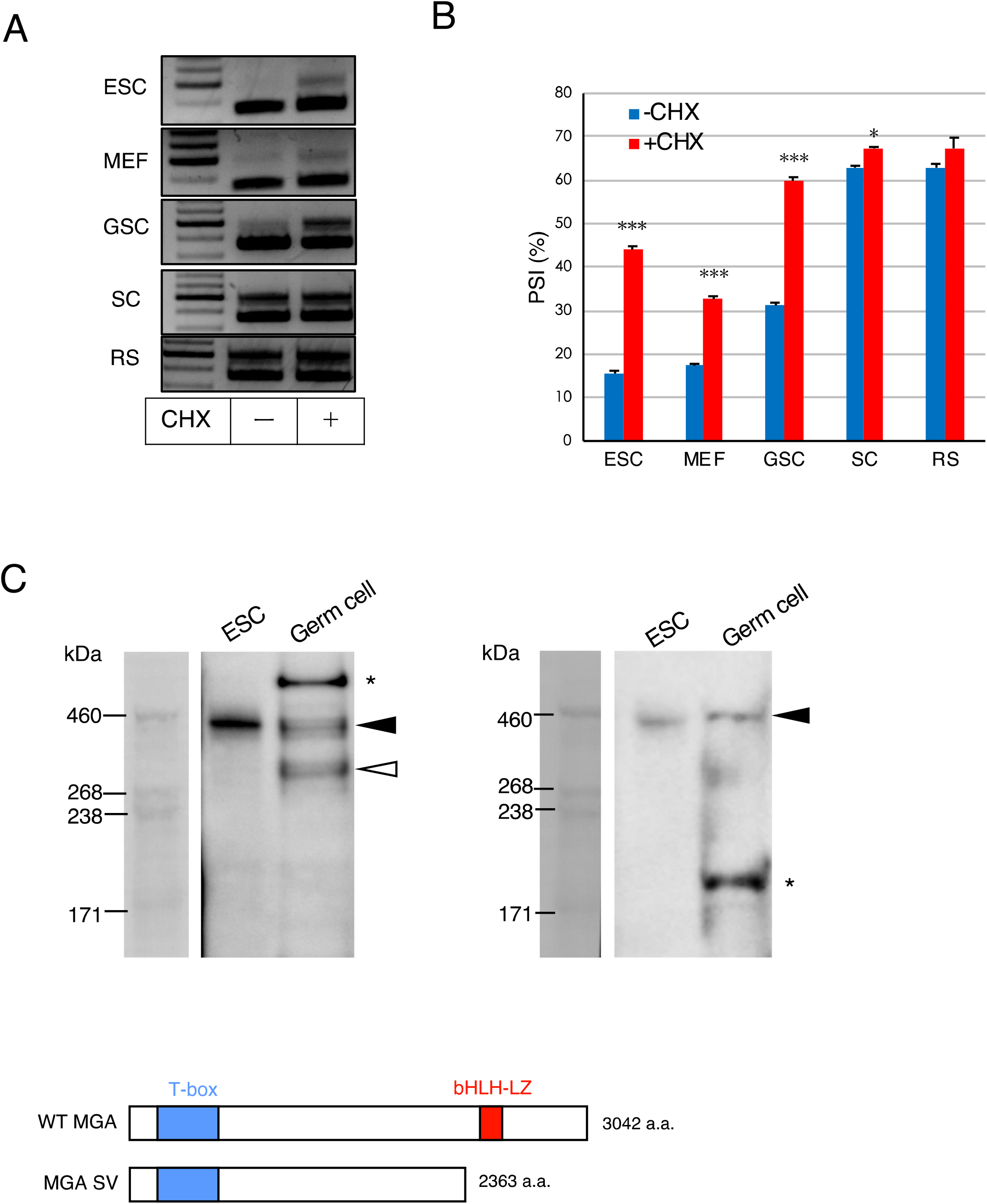
Generation of PTC-containing *Mga* SV mRNA leads to production of anomalous MGA protein in spermatocytes because of a low PTC-dependent NMD background. (A) High expression of *Mga* SV in spermatocytes and round spermatids was not further elevated by suppressing NMD activity. Five different cell types were treated with or without CHX for 3 h and total RNAs prepared from them were used to perform semi-quantitative PCR after conversion to cDNAs. (B) Quantification of wildtype and splice variant *Mga* mRNAs in non-germ and germ cells treated with or without CHX. cDNAs used in A were subjected to qPCR to accurately compare their amounts of wildtype and splice variant *Mga* mRNAs. Data are shown as the percent spliced in index (PSI; ratio of transcripts with exon inclusion versus total transcripts). Data represent the mean ± standard deviation of three independent experiments. The Student’s t-test was conducted to examine statistical significance. \**P*<0.05; \*\*\**P*<0.001 (C) Western blot analyses of MGA protein using nuclear extracts from ESCs and a mixture of spermatocytes and round spermatids. Antibodies that recognized amino-terminal (left panel) and carboxy-terminal (right panel) portions of MGA were used. Solid and open arrowheads indicate wildtype and anomalous MGA, respectively. *indicates a non-specific band.

### Dominant negative effect of carboxy-terminally truncated MGA on PRC1.6

To explore the possible function of MGA SV, we first examined the ability of the protein to interact with other PRC1.6 components by coimmunoprecipitation analyses (Fig. 5A). These analyses revealed no noticeable difference in the efficiency of the interaction with endogenous PCGF6, L3MBTL2, HP1γ, and RING1B in HEK293FT cells between flag-tagged wildtype MGA and its derivative, i.e., MGA SV, which were forcedly produced by transient transfection. Our data also demonstrated that both types of MGA proteins did not bind to SUZ12, a component of PRC2. In addition, we confirmed that wildtype MGA, but not MGA SV, interacted with MAX efficiently as expected. Next, we examined the effect of forced production of MGA SV on interactions of PRC1.6 components with genomic DNA. To this end, we also used the HEK293FT cell line. The advantage of using the HEK293FT cell line was complete disruption of *Mga* loci to eliminate the contribution of endogenous MGA in these cells without affecting their viability. In fact, we generated *Mga-KO* HEK293FT cells using the CRISPR-Cas9 system (Fig. S4) and used them to examine the interaction of overexpressed wildtype and anomalous MGAs with the promoters of PRC1.6 target genes (*CCND2, CDIP*, and *CNTD1*) whose repression is crucially dependent on the bHLHZ domain of MGA in these cells. These analyses revealed that wildtype MGA bound much more efficiently to all three gene promoters than MGA SV in *Mga*-null HEK293FT cells (Fig. 5B). At present, we do not know why the interaction scores obtained with MGA SV were not sufficiently low enough to judge as background. However, it is possible that these data represent binding of MGA SV to DNA using its T-box domain and/or other components of PRC1.6., i.e., E2F6/DP1 and L3MBTL2, which can lead to direct and indirect binding of the complex to DNA, respectively. Our analyses also revealed that binding of MAX, RING1B, and PCGF6 to these gene promoters was significantly deteriorated by forced expression of *Mga* SV, but not wildtype *Mga* (Fig. 5C), which suggested that MGA SV functioned as a dominant negative regulator of PRC1.6 binding to its target sites.

**Figure 5.**
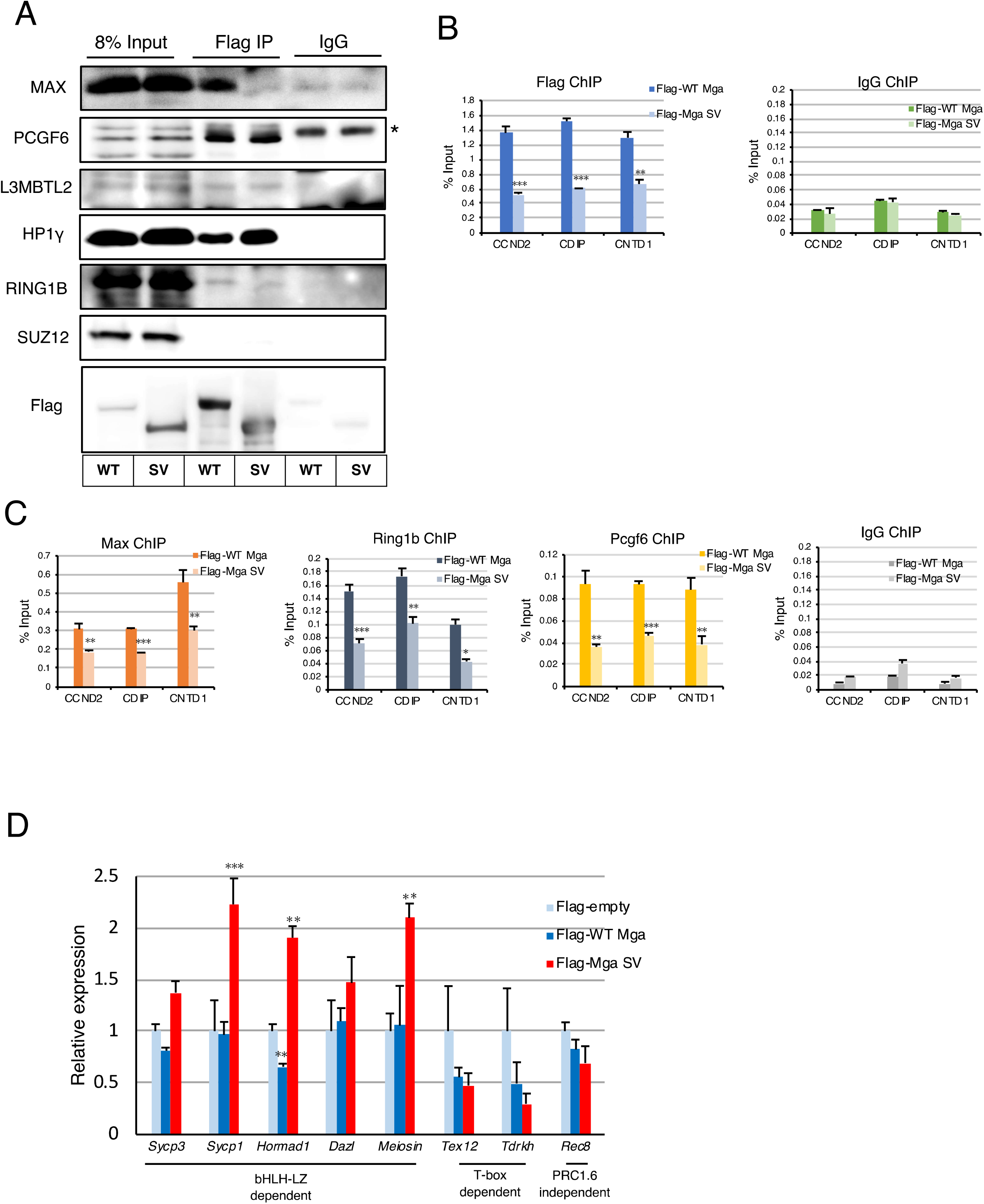
Potential of MGA SV as a dominant negative regulator of PRC1.6. (A) Coimmunoprecipitation analyses of wildtype and anomalous MGAs. Expression vectors for Flag-tagged wildtype and anomalous MGAs were transiently introduced individually into HEK293FT cells by transfection. Coimmunoprecipatations were performed with an anti-Flag-tag antibody using nuclear extracts prepared from the transfected cells. Coimmunoprecipitated proteins were used to examine the presence or absence of RING1B and SUZ12 as well as specific components of PRC1.6 (PCGF6, L3MBTL2, and HP1γ). *indicates signals of the immunoglobulin heavy chain used for immunoprecipitation. (B) ChIP analyses of PRC1.6-target genes in *Mga*-null HEK293FT cells producing Flag-tagged wildtype or anomalous MGA transiently with the anti-Flag-tag antibody. Control IgG was used as a negative control. Data represent the mean ± standard deviation of three independent experiments. The Student’s t-test was conducted to examine statistical significance. \*\**P*<0.01; \*\*\**P*<0.001 (C) ChIP analyses of PRC1.6-target genes in *Mga*-null HEK293FT cells producing Flag-tagged wildtype or anomalous MGAs transiently with antibodies against MAX, RING1B, or PCGF6. Control IgG was used as a negative control. Data represent the mean ± standard deviation of three independent experiments. Data were subjected to statistical analyses as described in B. \*\**P*<0.01; \*\*\**P*<0.001 (D) Forced expression of *Mga* SV in mouse ESCs was accompanied by significant activation of meiosis-related genes. Expression vectors to produce Flag-tagged wildtype and anomalous MGAs were transiently xintroduced individually into mouse ESCs by transfection. RNAs prepared from the transfected cells were used to quantify expression levels of meiosis-related genes. Data represent the mean ± standard deviation of three independent experiments. The Student’s t-test was conducted to examine statistical significance. \*\**P*<0.01; \*\*\**P*><0.001

Next, we used mouse ESCs in which PRC1.6 functions as a blocker of ectopic meiosis. We transiently introduced expression vectors for wildtype *Mga* and *Mga* SV individually and then examined alterations in the expression levels of meiosis-related genes. Our analyses revealed that some meiosis-related genes (*Sycp3, Sycp1, Hormad1, Dazl*, and *Meiosin*) indeed showed significant elevation in their expression levels after overexpression of *Mga* SV, but not wild-type*Mga*. However, other PRC1.6 target genes (*Tex12* and *Tdrkh*) were not activated, but further repressed by forced expression of *Mga* SV and wildtype *Mga*, while the *Rec8* gene, a meiosis-related gene that is not subjected to PRC1.6-dependent regulation, did not show appreciable alterations in its expression levels after overexpression of wildtype *Mga* or *Mga* SV (Fig. 5D). Because MGA SV lacks the bHLHZ domain, but carries an intact T-box domain, these data were in accordance with our recent observation that repression of *Tex12* and *Tdrkh* genes is dependent on the integrity of the T-box domain of MGA, while the bHLHZ domain of MGA is crucially involved in repressing the expression of *Sycp3, Sycp1, Hormad1, Dazl*, and *Meiosin* genes (34). Next, we investigated physiological alterations in the expression levels of meiosis-related genes during the conversion of spermatogonia to preleptone spermatocytes in male germ cells using publicly reported RNA sequence data. We found that meiosis-related genes whose repression was crucially dependent on the bHLHZ domain (*Sycp3* and *Sycp1*) profoundly elevated their expression levels during this conversion, whereas those primarily subjected to T-box-dependent repression, such as *Tdrkh* and *Tex12*, were much less significantly activated (Fig. S5). These results suggested that de-repression of a subset of meiosis-related genes by forced expression of *Mga* SV in ESCs faithfully recapitulated the physiological activation profile of meiosis-related genes during meiotic onset.

## Discussion

While the Stra8/Meiosin complex constitutes a positive feedback loop to promote meiotic onset in germ cells (2), PRC1.6 has the opposite activity (16, 19–21). Therefore, PRC1.6 needs to be inactivated at the timing of meiotic onset to produce a background that allows the Stra8/Meiosin complex to function efficiently and promote meiotic onset. In accordance with this notion, we have previously demonstrated that protein levels of MAX, which constitutes the core of PRC1.6 with MGA, decrease significantly immediately prior to the onset of meiosis in germ cells (19). In this study, we demonstrated the existence of an additional mechanism to alleviate the repressing activity of PRC1.6 for meiotic onset. Indeed, we demonstrated that *Mga* splice variant mRNA is specifically present in meiotic spermatocytes and postmeiotic round spermatids, which is translated into an MGA protein lacking the bHLHZ domain and functions as a dominant negative regulator of PRC1.6.

Thus, we have demonstrated previously (19) and in this study that at least two molecular mechanisms that target either Max or Mga operate in germ cells independently to impede the functions of PRC1. However, overlap between germ cells with reduced MAX protein levels and those positive for *Mga* SV mRNA is evident only up to the zygotene stage of meiotic prophase I in germ cells. Indeed, MAX protein levels increased significantly around the pachytene stage, while *Mga* SV mRNA was present during all stages of meiotic cell division and its presence was persistent at least up to the round spermatid stage. Thus, inactivation of PRC1.6 after the zygotene stage may be solely dependent on the dominant negative function of MGA SV generated from translation of *Mga* SV mRNA, although we do not eliminate the possibility that other unknown molecular mechanisms impede PRC1.6 in parallel.

MGA has two distinct DNA-binding domains termed T-box and bHLHZ domains that are independent and dependent on the interaction with MAX to bind to DNA, respectively (16, 35). However, it was recently demonstrated by an unbiased *de novo* sequence motif search that, unlike the E-box sequence that is recognized by the MGA/MAX complex via their bHLHZ domains, the T-box motif was not identified as a binding motif of PRC1.6 in chromatin without homozygous disruption of the *E2f6* gene in ESCs (16). Moreover, our recent study demonstrated that deletion of bHLHZ domain of MGA, in ESCs is accompanied by derepression of numerous meiosis-related genes that play pivotal roles in meiotic progression, such as *Meiosin* and *Sycp3*. However, only a few genes that have been demonstrated to be crucially involved in meiosis are significantly activated by deletion of the T-box domain (34). Although these data were obtained from studies using ESCs that bear the potential for ectopic meiosis, these studies strongly suggest that the bHLHZ domain has a much more predominant role in suppressing meiosis than the T-box domain. Therefore, we consider that production of the dominant negative MGA protein lacking the bHLHZ domain via alternative splicing is a reasonable strategy for meiotic germ cells to acquire high expression levels of biologically important meiosis-related genes to achieve initiation and progression of meiosis. Consistent with this assumption, we also noted that meiosis-related genes whose repression is strongly dependent on bHLH-LZ, such as *Sycp1* and *Sycp3*, dramatically elevated their expression levels during physiological transition from mitotic to meiotic cell division of male germ cells, whereas those primarily subjected to T-box-dependent repression, such as *Tdrkh* and *Tex12*, were not profoundly activated.

Unusual mRNAs such as those with a PTC including *Mga* SV are usually degraded rapidly by NMD (36). However, PTC-mediated NMD, but not long 3’-UTR-dependent NMD, is defective in spermatocytes and round spermatids. Therefore, conditional knockout of the *Upf2* gene, which is essential for NMD in these cells, is not associated with further accumulation of PTC-containing transcripts (32, 37). As the molecular mechanisms underlying this weak NMD activity in spermatocytes and round spermatids, *Upf3a* and *Upf3b* genes involved in the repression and activation of NMD, respectively, are transcriptionally up- and down-regulated in these germ cells. Moreover, conditional knockout of *Upf3a* in *Stra8*-positive meiotic germ cells is accompanied by substantial reductions in the levels of PTC-containing mRNAs in spermatocytes (38). Although we do not assume that the defective background of PTC-dependent NMD in meiotic spermatocytes and round spermatids is exclusive for production of anomalous MGA, to our knowledge, this is the first example that exquisitely uses this specialized condition for an obvious biological phenomenon.

In summary, our computational search identified a potential exon within the *Mga* gene and our subsequent analyses revealed that the sequence was incorporated into wildtype mRNA specifically in meiotic germ cells and round spermatids by alternative splicing. Moreover, our data revealed that these male germ cells produced anomalous MGA protein that may alleviate the repressing activity of PRC1.6 for meiotic onset as a dominant negative regulator by exquisitely using the inefficient background of PTC-dependent NMD in these cells.

## Materials and methods

### Animal ethics

This study was carried out under strict accordance with international and institutional guidelines. The protocol was approved by the Institutional Review Board on the Ethics of Animal Experiments of Saitama Medical University (permission numbers 2579 and 3032).

### Isolation of germ cells

To isolate Thy1^+^ and c-Kit^+^ spermatogonia, testes from C57BL/6 mouse pups at postnatal days 5–8 were collected and then minced after removal of the tunica albuginea membrane. After two consecutive enzymatic digestions using collagenase type I and trypsin along with DNase I, cells were recovered as floating cells in gelatin-coated tissue culture plates to enrich the germ cell population. The recovered cells were then subjected to magnetic activated cell sorting to isolate CD117 (c-Kit)^+^ cells followed by isolation of CD90 (Thy1)^+^ cells using the same method.

Spermatocytes and round spermatids were isolated from testes of 8–12-week-old adult C57BL/6 mice. After conducting the procedure described above, the testicular cell suspension was subjected to Percoll density gradient centrifugation and spermatocyte- and round spermatid-enriched fractions were isolated as fractions with 30% and 22% Percoll densities, respectively, by inspecting their nuclear morphology with an aid of Hoechst staining.

### RT-PCR and qPCR Analyses

Total RNAs and cDNAs were prepared as described previously (19). RT-PCR was performed using TaKaRa Ex Taq Hot Start Version (Takara) or PrimeSTAR Max DNA Polymerase (Takara). PCR products were subjected to agarose gel electrophoresis. TaqMan and SYBR Green-based qPCRs were performed using the StepOnePlus™ Real-Time PCR System (Applied Biosystems). To identify *Mga* variant mRNA by qPCR, one of the primers was set within a putative exon sequence (exon 19a). All samples were tested in triplicate and the results were normalized to *Gapdh* expression levels. Primer sequences and TaqMan probes are listed in Table 1.

### *In situ* hybridization for wildtype and splice variant *Mga* mRNAs

Formalin-fixed, paraffin-embedded sections of the testis and epididymis of adult ICR mice were used for BaseScope *in situ* hybridization assays employing a BaseScope Red reagent kit (Advanced Cell Diagnostics) (25) to detect wildtype and splice variant *Mga* mRNAs. The sections were counterstained with hematoxylin.

### Cell Culture

EBRTcH3 ESCs (39) were cultured in Glasgow minimum essential medium (Sigma) containing 10% fetal bovine serum and leukemia inhibitory factor (1000 U/ml). Mouse GSCs (kindly provided by Dr. Takashi Shinohara, Kyoto University, Japan) were cultured on mitomycin C-treated MEFs as described by Kanatsu-Shinohara et al. (40). HEK293FT cells were cultured in Dulbecco’s modified Eagle’s medium containing 10% fetal bovine serum.

### Cycloheximide treatment

ESCs, MEFs, and GSCs were treated with CHX (100 μg/ml) for 3 h. Spermatocytes and spermatids isolated from 8-week-old C57BL/6 mice were suspended in αMEM with 10% knockout serum replacement (Thermo Fisher Scientific) and 10 μg/ml GDNF (Funaskoshi), and then treated with or without CHX (100 μg/ml) for 3 h.

### Expression vector construction

To construct expression vectors for wildtype MGA and MGA SV, corresponding cDNAs recovered by PCR were individually introduced into the pCAG-IRES-Puro eukaryotic expression vector (41) using in-fusion technology after subcloning oligonucleotides carrying initiating methionine codon followed by a Flag-tag sequence.

### Generation of homozygous *Mga*-knockout HEK293FT cells

The CRISPR/Cas9 system was used to establish *Mga*-null HEK293FT cells. Oligonucleotide pairs carrying the same sequence used by Stielow et al. (16) were inserted into the pX330-U6-Chimeric_BB-CBh-hSpCas9 vector. The vector was then introduced into HEK293FT cells together with pCAG-mycGFP-IN by cotransfection using Lipofectamine 2000 (Invitrogen, Life Technologies). Subsequently, GFP-positive cells were collected by fluorescence-activated cell sorting and seeded a 96-well tissue culture plate with one cell per each well. Genomic DNAs prepared from individual cell colonies were used to identify clones in which the *Mga* gene was homozygously disrupted.

### Nuclear extract preparation, and coimmunoprecipitation and western blot analyses

Nuclei were prepared from 3×10^7^ HEK293FT cells by lysing the cell membrane under hypotonic conditions followed by low speed centrifugation (1,000 rpm). Then, nuclei were resuspended in hypertonic buffer containing 300 mM NaCl and placed on ice for 15 min. A nuclear extract was prepared as the supernatant after moderately high speed centrifugation (10,000 rpm) as described by Uranishi et al. (42). For coimmunoprecipitation analyses, an anti-Flag antibody or normal mouse IgG were added to the nuclear extract together with Dynabeads bearing anti-Mouse IgG (Thermo Fisher Scientific). After incubation for 1 h with gradual rotation at 4°C, proteins bound to beads were eluted with sample buffer after extensive washing with PBS containing 0.1% BSA, heated, and then resolved by SDS-PAGE. Separated proteins were electrophoretically transferred to PVDF membranes. The membranes were blocked in PBST containing 5% dry skim milk and subjected to western blot analyses with primary antibodies followed by appropriate secondary antibodies conjugated with horseradish peroxidase. The antibodies are listed in Table 2.

### ChIP-qPCR Analysis

Suspended cells were fixed with 1% paraformaldehyde in PBS for 10 min at room temperature and then washed extensively with PBS. The cells were resuspended in cell lysis buffer (10 mM Tris-HCl, pH 8.1, 10 mM NaCl, 1.5 mM MgCl_2_, and 0.5% Igepal-CA630) with a protease inhibitor cocktail and placed on ice for 15 min. After centrifugation, pellets were resuspended in nuclear lysis buffer (50 mM Tris-HCl, pH 8.1, 5 mM EDTA, and 1% SDS) with a protease inhibitor cocktail and then sonicated. After centrifugation, the supernatant was subjected to an immunoreaction with a specific antibody pre-conjugated with magnetic protein A beads (Millipore) overnight at 4°C. The beads were washed twice with TE buffer after consecutive washes with low salt wash buffer (0.1% SDS, 1% Triton X-100, 2 mM EDTA, 20 mM Tris-HCl, pH 8.1, and 150 mM NaCl), high salt wash buffer (0.1% SDS, 1% Triton X-100, 2 mM EDTA, 20 mM Tris-HCl, pH 8.1, and 500 mM NaCl), and LiCl wash buffer (1% Igepal-CA630, 1% deoxycholate, 1 mM EDTA, 10 mM Tris-HCl, pH 8.1, and 250 mM LiCl). Crosslinks were cleaved by heat treatment. Genomic DNAs associated with beads were recovered using a QIAquick PCR Purification Kit (Qiagen) and then used to quantify their amounts by qPCR.

### Publicly Reported RNA Sequencing Data Analyses

To compare alterations in the frequency and predominant types of alternative splicing during spermatogenesis, RNA sequence data reported by Lin et al. (43) (GSE75826) were retrieved. Likewise, to assess the dynamics of alternative splicing during conversion of neural progenitor cells to neuronal cells and differentiation of MSCs into osteoblasts, RNA sequence data reported by Liu et al. (44) (GSE96950) and Shao et al. (45) (GSE112694) were retrieved, respectively. To construct the Sashimi plots shown in Figure 3B and C, RNA sequence data reported by Lin et al. (43) (GSE75826) and Sangrith et al. (46) (E-MTAB-4616) were used, respectively. Retrieved fastq files were mapped to mouse genome version mm9 with HISAT2 version 2.1.0 and analyzed using the Tuxedo protocol (47).

### Alternative Splicing Analyses

Aligned bam files were analyzed using MISO version 0.5.4. The threshold to be judged as differentially expressed between two different cell types was set to 10 in Bays factor.

Sashimi plots of RNA sequence reads were generated by Integrative Genomics Viewer.

## Supporting information

Supplemental items

## Acknowledgments

The authors thank Dr. Robert N. Eisenman for his provision of Mga cDNA and helpful discussions. The authors are also indebted to Drs. Kishore Jaganathan and Kyle Kai-How Farh for their attentive instructions for the use of the SpliceAI deep learning tool. We thank Mitchell Arico from Edanz Group (https://en-author-services.edanzgroup.com/ac) for editing a draft of this manuscript. This work was supported in part by the Ministry of Education, Culture, Sports, Science and Technology (MEXT), Japan. K.U. and A.O. are recipients of grants from the Japan Society for the Promotion of Science (JSPS) KAKENHI (grant numbers 20K16147 and 19H03426, respectively). This work was also supported in part by the Research Fellowship for Young Scientists (DC2) from JSPS to Y.K (grant number 20G10148).

## Author Contributions

Y.K., K.U., A.S., and A.O. designed the study; Y.K., K.U., and A.S. performed experiments; M.H., and M.N. contributed reagents/analytical tools; Y.K., K.U., M.H., M.N., and A.S. analyzed data; Y.K. and A.O. wrote the manuscript.

The authors declare no competing interest.

