## Supplemental items for "Identification of germ cell-specific *Mga* variant mRNA that promotes meiotic entry via impediment of a non-canonical PRC1": Kitamura et al. supplemental Figures and Tables 8-31-2020.pdf

for manuscript entitled

The file contains five Figures and two Tables

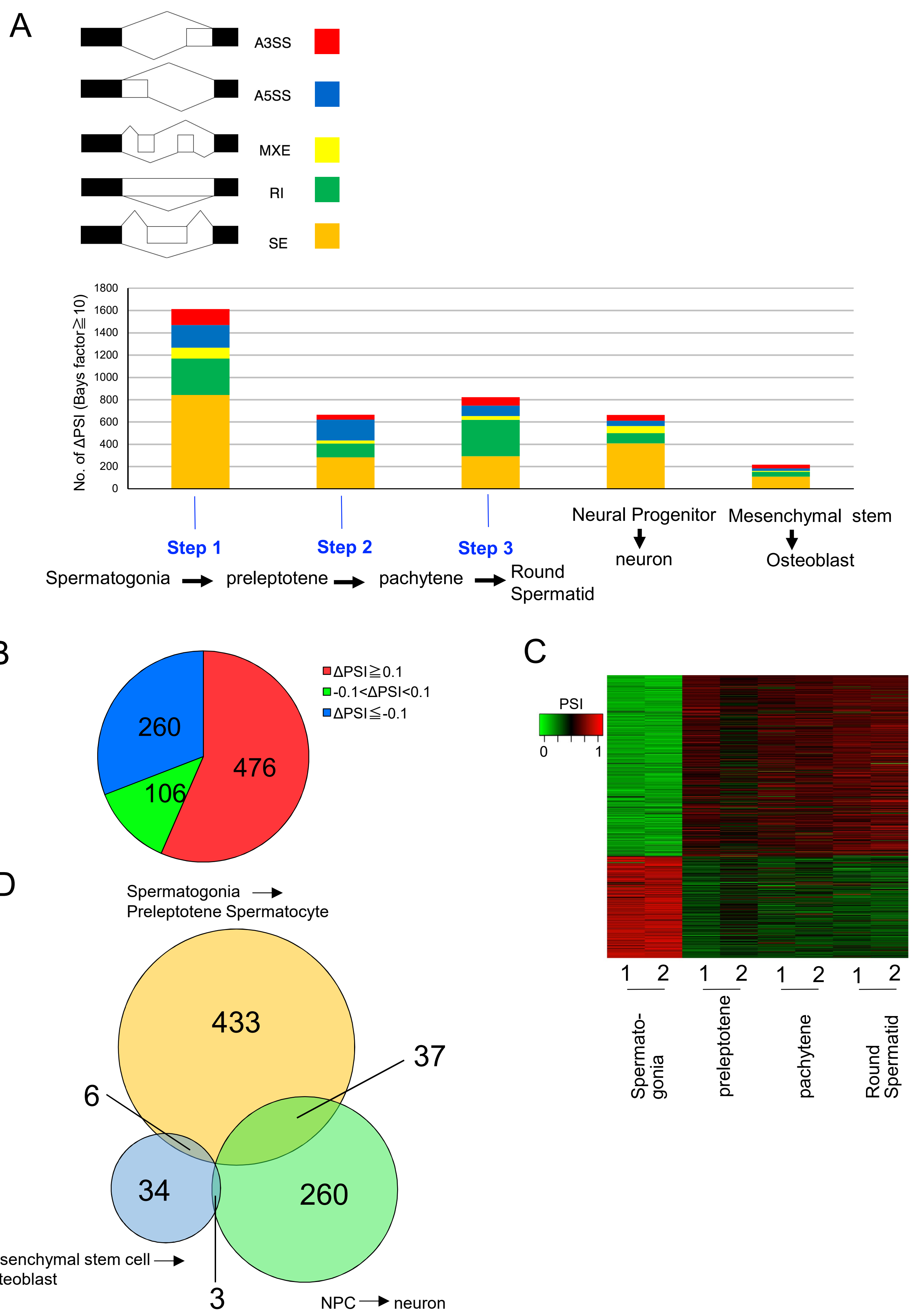

A
A3SS
A5SS
MXE
RI
SE
No. of ΔPSI (Bays factor≧10)
1800
1600
1400
1200
1000
800
600
400
200
0
Step 1
Step 2
Step 3
Spermatogonia
preleptotene
pachytene
Round Spermatid
Neural Progenitor
neuron
Mesenchymal stem
Osteoblast
B
ΔPSI≧0.1
-0.1<ΔPSI<0.1
ΔPSI≦-0.1
260
476
106
C
PSI
0
1
1 2
1 2
1 2
1 2
Spermato-gonia
preleptotene
pachytene
Round Spermatid
D
Spermatogonia
Preleptotene Spermatocyte
433
37
6
34
260
3
Mesenchymal stem cell
Osteoblast
NPC
neuron

**Figure S1. SE type is the most predominant alternative splicing during meiotic onset in germ cells** (A) Comparisons of frequency and the types of alternative splicing among the three different transitions in stage of spermatogenesis. Five different types of alternative splicing are schematically shown at the top. Publicly reported RNA sequence data were used to obtain the PSI in the four different germ cell types indicated in the schema. A bar graph was constructed after calculating changes in the PSI (ΔPSI) at steps 1–3 individually. Data from differentiations of neural progenitor cells and mesenchymal stem cells are also shown as references. A3SS: alternative 3' splice site; A5SS: alternative 5' splice site; MXE: mutually exclusive exon; RI: retained intron; SE: skipping exon (B) Classification of SE type alternative splicing into three subgroups according to the range of ΔPSI. (C) Transcripts with an increased or decreased PSI during meiotic onset were maintained at least up to round spermatids. Genes with PSIs that were significantly increased or decreased at the stage corresponding to step 1 in A were selected and their PSIs in spermatogonia, preleptene spermatocytes, pachytene spermatocytes, and round spermatids were plotted. Data were retrieved from two independent experiments (1 and 2) conducted by Lin et al. (43). (D) Venn diagram showing comparisons of genes with ΔPSIs that is equal or larger than 0.1 in the differentiation of spermatogonia, neural progenitor, or mesenchymal stem cells. No genes that showed 0.1 or larger ΔPSI values upon differentiation were shared among the three different cell types.

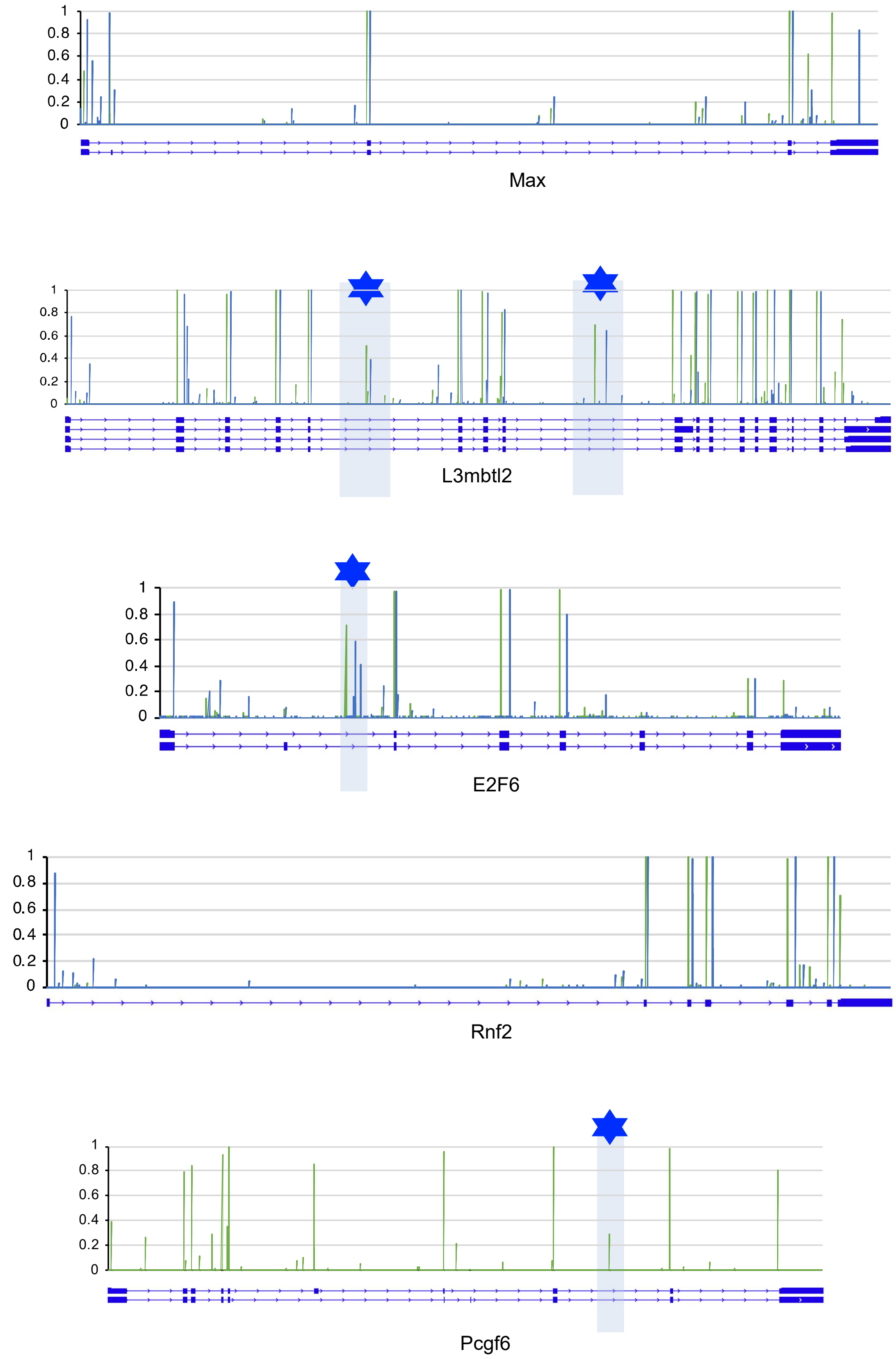

**Figure S2. Search for potential exons within genes encoding a PRC1.6 component by SpliceAI deep learning.** Sequences from pre-mRNA transcripts of genes encoding a PRC1.6 component (*Max, L3mbtl2, E2f6, Rnf2*, and *Pcgf6*) were subjected to the analyses of SpliceAI deep learning. Scores as the splice acceptor and donor are shown as green and blue bars, respectively. Blue asterisks indicate regions with a set of high scores for the splice acceptor and donor determined by SpliceAI except for known exons.

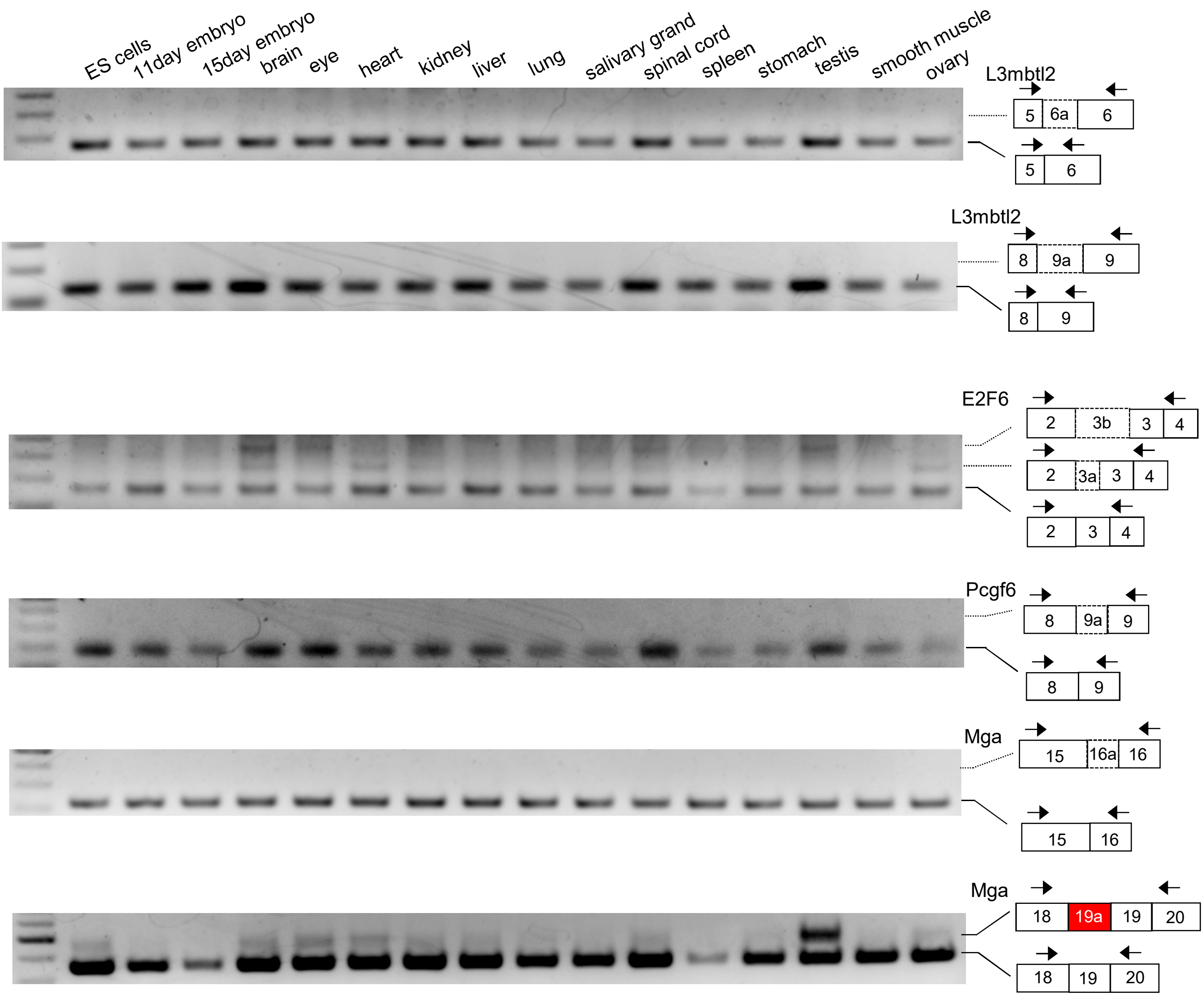

**Figure S3. RT-PCR analyses of the regions identified as putative exons by SpliceAI.** RT-PCR analyses were conducted using 16 different mRNAs with respect to the regions suggested as putative exons by SpliceAI. Several faint and/or smear bands obtained by analyses of the *E2f6* gene in addition to the band corresponding to the wildtype were found to be irrelevant bands by sequencing PCR products.

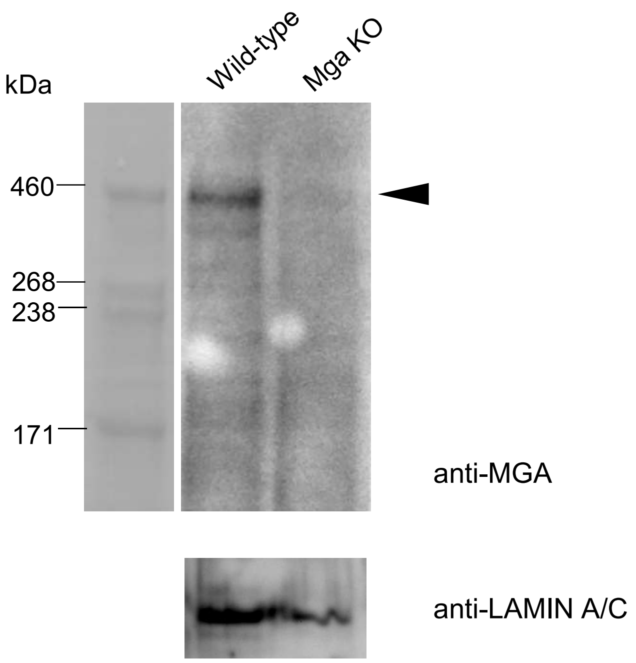

**Figure S4. Confirmation of the lack of MGA in *Mga*-knockout HEK293FT cells by western blot analyses.** MGA and internal control LAMIN A/C proteins were detected by western blot analyses of nuclear extracts from wildtype and *Mga*-null HEK293FT cells. Homozygous knockout of the *Mga* gene in HEK293FT cells was conducted by CRISPR-Cas9-mediated genomic manipulation targeting the region around the 3′-end of exon 3 of the *Mga* gene using oligonucleotide sequences described by Stielow et al. (16). Generated *Mga*-null HEF293FT cells were identical to those generated by Stielow et al. (16) at the single nucleotide sequencing level, i.e., 73 and 55 bp deletion in the 3′-end of exon 3 and 5′-end of intron 3, respectively, causing abnormal splicing and a frameshift.

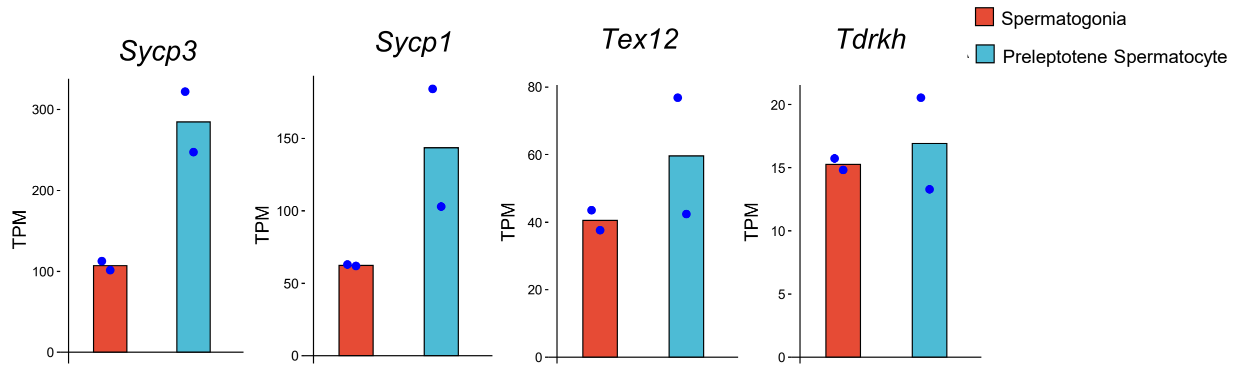

**Figure S5. Expression levels of meiosis-related genes during meiotic onset in publicly reported RNA sequence data.** Expression data of meiosis-related genes that are primarily subjected to regulation by bHLHZ (*Sycp3* and *Sycp1*) or the T-box domain (*Tex12* and *Tdrkh*) of MGA in spermatogonia and preleptotene spermatocytes were extracted from publicly reported RNA sequence data by Lin et al. (43) and shown as a bar graph. Data are shown as the mean of two independent experiments in which each dot represents the value from an individual experiment.

### Table S1 Oligonucleotides and TaqMan Probes

#### Oligonucleotides

##### RT-PCR

| | | |
|---|---|---|
| L3mbtl2 Exon5-6 | forward | 5'-ACTGGGGCAAGTTCCTGAAG-3' |
| | reverse | 5'-TGCCTGGATGACAGTGGCGAT-3' |
| L3mbtl2 Exon8-9 | forward | 5'-GAAGAGCTACCTCATGAAGCGG-3' |
| | reverse | 5'-CACCTGAGTCTTGTCTACAACCTC-3' |
| E2F6 Exon 2-4 | forward | 5'-CTTCTAGCCAGGTGTGGTGG-3' |
| | reverse | 5'-TACCAGTGACACATCAAACCGG-3' |
| Pcgf6 Exon8-9 | forward | 5'-CCATTGGAAAAGAAGTTTGTGCGTG-3' |
| | reverse | 5'-GCTGTATCACCTATTGCACGTCG-3' |
| Mga Exon 15-16 | forward | 5'-GGTGACCACACCTACTTCATCACTG -3' |
| | reverse | 5'-TGTCCCTGAAGCTGTGGGTT-3' |
| Mga Exon18-19 | forward | 5'-GAGGATGAGGAAGATGAGAAAACTGA-3' |
| | reverse | 5'-TGTCCGTCGGTAATATGCAA-3' |

##### ChIP-qPCR

| | | |
|---|---|---|
| CCND2 | forward | 5'-CGCCACCAGATCGTATCTCCTGTAA-3' |
| | reverse | 5'-CCTCACTCGCCAGGCTTTCT -3' |
| CDIP | forward | 5'-CAGCCTCGTGTACATTGGGCA-3' |
| | reverse | 5'-GAGGCGATTTGGCCTAGAGCT-3' |
| CNTD1 | forward | 5'-GTAGGACCTTCTGCCACTGGG-3' |
| | reverse | 5'-GAGCTGGTGACCCTCTGGATTCT-3' |

##### qPCR

| | | |
|---|---|---|
| Meiosin | forward | 5'-CATTGACATGACCAAGGCCTTGC-3' |
| | reverse | 5'-TGGAGGGAGTGGAGTGTTGCT -3' |
| Tex12 | forward | 5'-GAGAAGGATTTGAGCGATATGAGCAAGG-3' |
| | reverse | 5'-CTGTAAACCTCTGCTTCAGGAACTCTC -3' |
| Tdrkh | forward | 5'-TTCTGGTGCCCAGAGCAGTC-3' |
| | reverse | 5'-GGCTGCGGGAACCAATGATTTG -3' |

##### CRISPRE/Cas9

| | | |
|---|---|---|
| hMGA Exon 3 | forward | 5'-CACCGCATCTGGAAAGGTACTCCCA-3' |
| | reverse | 5'-AAACTGGGAGTACCTTTCCAGATGC-3' |
| hMGA intron 3 | forward | 5'-CACCG TCATACTTGAATTGTATAC-3' |
| | reverse | 5'-AAACGTATACAATTCAAGTATGAC-3' |

**Genotyping for MGA-KO HEC293FT**

| | | |
|---|---|---|
| hMGA Exon3-intron3 | forward | 5'-GAAAGAGCCTCAGTGGAAATATCCTG-3' |
| | reverse | 5'-ATGAAAATTCCAGTAAGACCCGAAGAC-3' |

#### TaqMan probes used for qPCR

| Gene Symbol | Probe ID | |
|---|---|---|
| *Sycp1* | Mm01298009_m1 | |
| *Sycp3* | Mm00488519_m1 | |
| *Hormad1* | Mm00471448_m1 | |
| *Dazl* | Mm03053726_s1 | |
| *Rec8* | Mm00490939_m1 | |
| *Gapdh* | Mm99999915_g1 | |
| wild-type *Mga* | **Custom-made** | |
| | forward | 5'-GAAGACCACAGCAACTCACACAC-3' |
| | reverse | 5'-TTTTTCATCTGCAGAGATATGGCTA-3' |
| | probe | 5'-TCCTTCAAACAGCAGTGTC-3' |
| variant *Mga* | **Custom-made** | |
| | forward | 5'-GATTCCTGAGACAGTTTCCTAAGTGA -3' |
| | reverse | 5'-TTTTTCATCTGCAGAGATATGGCTA-3' |
| | probe | 5'-TTCAGTTACCTATTAAGGTGTC-3' |

### Table S2 Antibodies

**Primary Antibodies**

| Antigen | Manufacturer | Catalog No. | Usage | Remark |
|---|---|---|---|---|
| mouse MGA | abcam | ab214814 | WB | rabbit monoclonal |
| human MGA* | | | WB | rabbit polyclonal |
| mouse MAX | SANTA CRUZ | sc-197 | ChIP | rabbit polyclonal |
| human MAX | Proteintech | 10426-1-AP 1 | WB | rabbit polyclonal |
| human PCGF6 | abcam | ab192395 | WB | rabbit polyclonal, cross-reacts with mouse PCGF6 |
| mouse PCGF6 | abcam | ab200038 | ChIP | rabbit monoclonal |
| human L3MBTL2 | SANTA CRUZ | sc-365134 | WB | mouse monoclonal |
| human HP1γ | SANTA CRUZ | sc-398562 | WB | mouse monoclonal, cross-reacts with mouse HP1γ |
| human RING1B | Cell Signaling | #5694 | WB, ChIP | rabbit monoclonal l, cross-reacts with mouse RING1B |
| human SUZ12 | abcam | ab12073 | WB | rabbit polyclonal, cross-reacts with mouse SUZ12 |
| FLAG-tag | Sigma-Aldrich | F3165 | WB, ChIP, IP | mouse monoclonal |
| Normal Rabbit IgG | Cell Signaling | #2729 | ChIP | use as control IgG in ChIP experiments |
| Normal Mouse IgG1 | Cell Signaling | #5415 | ChIP, IP | use as control IgG in ChIP and IP experiments |

**Horseradish Peroxidase- Conjugated Secondary Antibodies**

| Antigen | Manufacturer | Catalog No. | Usage | Remark |
|---|---|---|---|---|
| rabbit IgG | Cell Signaling | #7074 | WB | goat polyclonal |
| mouse IgG | Cell Signaling | #7076 | WB | horse polyclonal |
| rabbit IgG | ROCKLAND | 18-8816-33 | WB | mouse monoclonal |
| mouse IgG | ROCKLAND | 18-8817-33 | WB | rat monoclonal |

*kindly provided by Dr. Bastial Stielow at Institute of Molecular Biology and Tumor Research, Philipps-University of Marburg in Germany
